# Robust self-organization of livestock pluripotent stem cells into post-gastrulation embryo models with advanced neuronal and mesodermal structures

**DOI:** 10.64898/2026.03.02.709113

**Authors:** Michelle Hauser, Peleg Berkowicz, Michael Namestnikov, Benjamin Dekel, Sharon Schlesinger, Iftach Nachman

**Affiliations:** Tel Aviv University; The Hebrew University

## Abstract

Mammalian body plan formation arises from the self-organization of pluripotent cells through conserved morphogenetic processes that are difficult to study in vivo. Stem cell–based embryo models (SEMs) offer accessible three-dimensional systems to investigate these events but are currently limited to mouse and human cells and largely recapitulate posterior embryonic structures. In addition, no in-vitro models exist for post-gastrulation development in ungulate species, whose early development differs from that of rodents and primates.

Here, we establish SEMs for two common ungulates, sheep and pig, using pluripotent stem cell–derived aggregates. We generate ovine and porcine gastruloids that recapitulate key features of gastrulation, including germ layer specification, symmetry breaking, and axial elongation. We further develop ovine trunk-like structures (oTLSs) that robustly model post-gastrulation trunk development, exhibiting sustained elongation, neuromesodermal progenitor maintenance, segmented somite formation, and a central neural tube–like axis. Time-resolved single-cell RNA sequencing combined with immunostaining reveals coordinated emergence of neural, mesodermal, and intermediate mesodermal lineages arranged along an anteroposterior axis. Notably, oTLSs generate dorsal neural derivatives, anterior neuronal populations, and renal primordia, representing an expansion in the lineage repertoire reported for existing trunk models.

Together, this work extends SEMs to livestock species and establishes a platform for comparative mammalian developmental studies, with potential applications in fundamental research, veterinary toxicology, and agricultural biotechnology.

## Introduction

Understanding how the mammalian body plan emerges from pluripotent cells requires experimental access to the coordinated, multi-lineage morphogenetic events that occur during development. While much of our current knowledge derives from rodent and human embryos, early development in large mammals such as sheep and pig follows distinct trajectories, including prolonged peri-implantation stages, with somitogenesis onset preceding implantation ^1,2^. These species occupy an important comparative position, bearing differences in embryonic geometry and occupying an intermediate position in developmental tempos between mouse and human ^3^. However, direct study of ungulate embryos remains technically challenging due to their in’utero development.

Stem cell–based embryo models (SEMs) have emerged as powerful in vitro systems to study early mammalian development under controlled conditions ^4^. Three-dimensional models such as gastruloids ^5, 6, 7, 8^ and trunk-like structures (TLSs), developed in mouse ^9, 10^ and human ^11, 12^ pluripotent stem cells, recapitulate important aspects of gastrulation and post-implantation morphogenesis, including symmetry breaking, germ layer specification, axial elongation, and early trunk patterning. These systems enable quantitative, perturbative, and time-resolved analyses that are difficult to achieve in vivo. Nevertheless, current SEMs are restricted to rodent and primate species and predominantly capture posterior embryonic programs, with limited generation of anterior neural and organ-associated lineages, with a recent exception that recapitulates more anterior structures by aggregating differently pre-treated cells ^13^. Recent advances in isolation and maintenance livestock ESCs open the door for in vitro modeling of ungulate species ^14, 15^

Using recently established embryonic disc-like PSC lines ^16^, we establish stem cell–based embryo models for two common ungulates, sheep and pig, using pluripotent stem cell–derived aggregates. By systematically deconstructing and adapting protocols used in mouse and human ESC-based systems, we develop reproducible protocols for generating ovine and porcine gastruloids that undergo symmetry breaking and axial elongation while exhibiting distinct dorso–ventral patterning biases. Building on the ovine gastruloid and by introducing ECM, we generate robust ovine trunk-like structures that sustain neuromesodermal progenitors, undergo prolonged axial elongation, and reproducibly form segmented somites flanking a central neural tube–like structure and develop neural crest, anterior neural tissues, kidney progenitors, and endothelial cells.

Using time-resolved single-cell RNA sequencing and immunostaining, we show that ovine trunk-like structures self-organize into a multi-lineage trunk architecture consistent with anteroposterior patterning inferred from HOX temporal dynamics and localized marker domains. These structures generate dorsal neural derivatives alongside paraxial and intermediate mesoderm populations, extending beyond the lineage repertoire of existing trunk models. Together, this work establishes ungulate SEMs as tractable platforms for comparative studies of mammalian post-implantation development and reveals the capacity of large-mammal pluripotent cells to self-organize into complex embryonic structures in vitro.

## Results

### Ungulate pluripotent stem cells self-organize into gastruloids

To assess the self-organizing capacity of ungulate pluripotent stem cells, we adapted established mouse and human gastruloid protocols ^17, 6^ to ovine and porcine embryonic disc-like stem cells, while adjusting for developmental tempo and the pluripotency state of the starting PSCs ^18^. Cells were maintained under AFX (ActivinA, FGF2, WNT inhibition by XAV939) pluripotency conditions and were transitioned to a gastrulation-competent state by withdrawal of XAV939 followed by transient WNT activation and subsequent aggregation in ultra–low-attachment U-bottom plates (Fig. 1A). Following aggregation, both ovine and porcine aggregates underwent rapid morphological reorganization, with symmetry breaking evident within 24h and progressive axial elongation over 72h (Fig. 1B).

**Figure 1.**
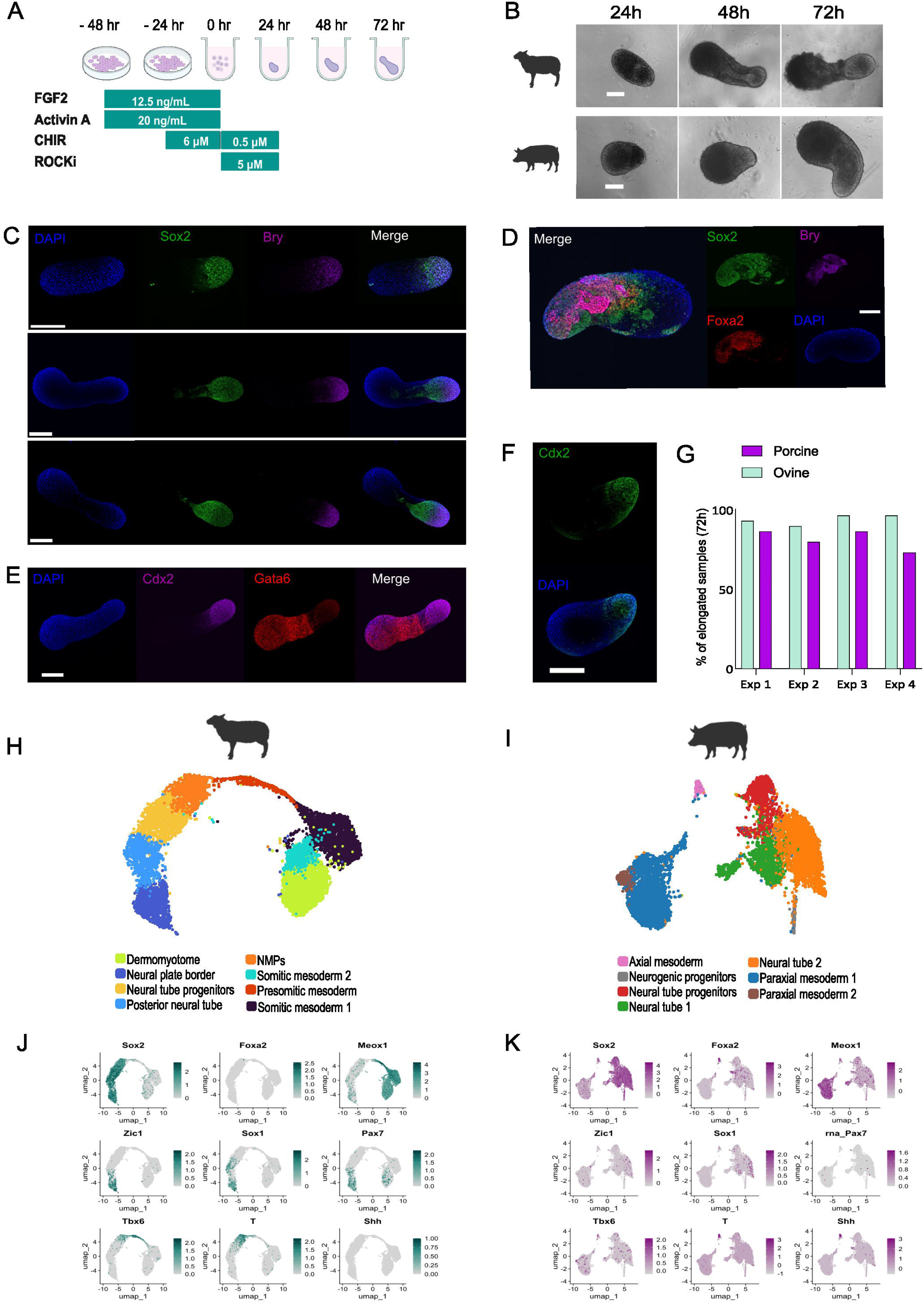
Ovine and porcine gastruloids display symmetry breaking, elongation and lineage segregation. (A) Graphical representation of protocol for the formation of ovine and porcine ESC-based gastruloids. (B) Representative images of ovine (top) and porcine (bottom) gastruloids at 24h, 48h and 72h. (C) Immunofluorescence-labeled ovine gastruloids at 24h, 48h and 72h, stained for early lineage markers Sox2 and Brachyury. (D) Representative porcine gastruloid at 72h stained for early lineage markers Bry, Sox2 and Foxa2. (E) 72h ovine gastruloid stained for posterior marker Cdx2 and anterior Gata6. (F) 72h porcine gastruloid stained for posterior marker Cdx2. (G) Efficiency of gastruloid protocol across 4 experimental repeats. N=120. (H-I) Integrated UMAP of scRNA-seq profiles from pools of 48□h and 72□h ovine (H) and porcine gastruloids (I). (J) Normalized expression of ovine gastruloid marker genes. (K) Normalized expression of porcine gastruloid marker genes. Scale bars 150 µm.

In ovine aggregates, immunofluorescence revealed polarized expression of Brachyury (Bry) and Sox2 at the posterior end, consistent with the emergence of a neuromesodermal progenitor (NMP)–like population. Over time, these domains segregated, with Sox2□ cells extending along the midline to form an elongated neural axis and Bry□ cells remaining confined posteriorly (Fig. 1C). Foxa2 expression was not detected in ovine aggregates. In porcine aggregates, Sox2 exhibited posterior polarization, but Bry was internally distributed and overlapped extensively with Foxa2 by 72h (Fig. 1D); this Bry and Foxa2 patterning is consistent with an axial mesoderm or mesendoderm–like population rather than a canonical NMP domain, ^19^. Consistent with early anteroposterior polarization, aggregates of both species showed posterior Cdx2 and opposing Gata6 expression within 24h of aggregation (Fig. 1E,F). Symmetry breaking occurred in the majority of aggregates, with 94% of ovine and 81% of porcine samples exhibiting polarized morphologies (Fig. 1G).

Ovine gastruloids displayed a dose-dependent elongation response to WNT pathway activation. Increasing concentrations of the WNT agonist CHIR99021 during pretreatment resulted in progressively elongated morphologies at 72h (Fig. S1A), with length-to-width measurements confirming a significant, concentration-dependent effect (n = 24; P < 0.005) (Fig. S1B).

To further characterize cellular composition, we performed single-cell RNA sequencing on ovine and porcine gastruloids collected at days 2 and 3 post-aggregation. In ovine gastruloids, UMAP analysis identified a prominent Bry□Sox2□ NMP-like population positioned between diverging neural and mesodermal trajectories. The mesodermal branch progressed toward paraxial and somitic states, while the neural lineage advanced toward a neural plate border–like population; no definitive endodermal population was detected (Fig. 1H). In contrast, porcine gastruloids lacked a comparable NMP-like population and instead contained a Bry□Foxa2□ axial mesoderm–like cluster alongside neural and mesodermal populations (Fig. 1I).

In both species, the somitic marker Tcf15 was expressed within mesodermal clusters, consistent with paraxial mesoderm identity; however, dorsal patterning differed markedly. In porcine gastruloids, Pax3 expression was confined to mesodermal clusters and absent from neural populations, whereas ovine gastruloids exhibited Pax3 expression across both neural and mesodermal lineages. This dorsal bias in ovine gastruloids was further supported by Pax7 expression, which was detected in ovine but not porcine datasets (Fig. 1J,K). These dorsoventral patterning differences correlated with the presence of NMPs in ovine gastruloids and a ventral axial mesoderm–skewed transcriptional profile in porcine gastruloids.

Despite these molecular differences, both ovine and porcine aggregates progressed through the defining morphogenetic stages of gastruloid development, including symmetry breaking, axial organization, elongation, and lineage segregation. Together, these findings demonstrate that ungulate pluripotent stem cells can generate bona fide gastruloid models in vitro and provide a foundation for studying early post-implantation development in large mammals.

### Ovine gastruloids develop into robust trunk-like structures upon ECM introduction

Encouraged by the evidence indicating the presence of NMPs and their derivatives in ovine gastruloids, we next examined whether the addition of extracellular matrix could promote higher-order tissue organization, as previously shown in mouse and human TLS systems ^20, 11^. We systematically refined parameters involved in TLS formation, including the timing of ECM addition, WNT activator concentration and timing, and the use of additional patterning cues (Fig. S2A–C). Introduction of Geltrex at 24h post-aggregation, immediately following symmetry breaking, proved critical for reproducible TLS morphogenesis. Withdrawal of the WNT inhibitor XAV939 prior to transient CHIR activation improved developmental outcomes, whereas inclusion of retinoic acid consistently resulted in aberrant morphologies and was therefore excluded from subsequent experiments. During optimization, we observed that early aggregate morphology was predictive of downstream developmental success. Specifically, increased asphericity at 24h post-aggregation strongly correlated with subsequent TLS formation, providing an early morphological criterion to identify aggregates with high developmental potential and serving as a morphological cue for the correct timing of ECM introduction ^21^ (Fig. S2D).

We next examined the influence of WNT activation strength on TLS morphogenesis. Consistent with earlier gastruloid experiments, increasing CHIR concentrations promoted greater axial elongation. A transient 6 µM CHIR pulse prior to aggregation yielded the most reproducible and well-organized structures characterized by elongated morphologies with segmented somite-like domains flanking a central axial neural tube–like region (Fig. S2E).

Ovine TLSs (oTLSs) exhibited a high degree of robustness across a wide range of initial aggregate sizes. Aggregates generated from 400 to 8,000 initial cells consistently underwent axial elongation and developed a well-organized central tube-like structure flanked by segmented somite-like domains (Fig. S3A). Across this 20-fold range in initial size, elongation dynamics and overall growth trajectories were comparable: oTLS elongation progressed until approximately day 6, after which structures reached a plateau. Aggregates generated from 1,000 cells or more maintained structural integrity beyond this point and did not exhibit collapse or regression through at least day 7 in culture (Fig. S3B,C). A starting size of 4,000 cells per aggregate was selected for subsequent experiments. This intermediate condition reproducibly generated well-defined somite-like segmentation and a clearly delineated neural tube–like structure while maintaining stable and interpretable morphologies at later developmental stages (days 7–8).

The optimized protocol (Fig. 2A) enabled the reliable and reproducible generation of oTLS; this success was not reproduced in porcine cells, and subsequent results deal exclusively with oTLSs. Across experiments, oTLSs consistently formed a central neural tube–like structure aligned along the midline and flanked bilaterally by segmented somite-like units (Fig. 2B). These structures continued to mature over time, exhibiting progressive elongation and sequential addition of somite-like segments through day 8 post-aggregation (Fig. 2C). oTLSs displayed an extended window of structural integrity: rather than collapsing, aggregates continued to elongate and grow until axial tissues became indescernable as discrete morphological units by approximately day 8. In addition to temporal stability, oTLS formation was highly reproducible within and across experimental replicates. Using the optimized protocol,over 90% of aggregates elongated along a single, well-defined axis (Fig. 2D). Furthermore,more than 80% of aggregates generated both clearly identifiable neural tube–like and segmenting somite-like structures (Fig. 2E). A minority of aggregates either failed to elongate or developed atypical morphologies, including dual elongation axes.

**Figure 2.**
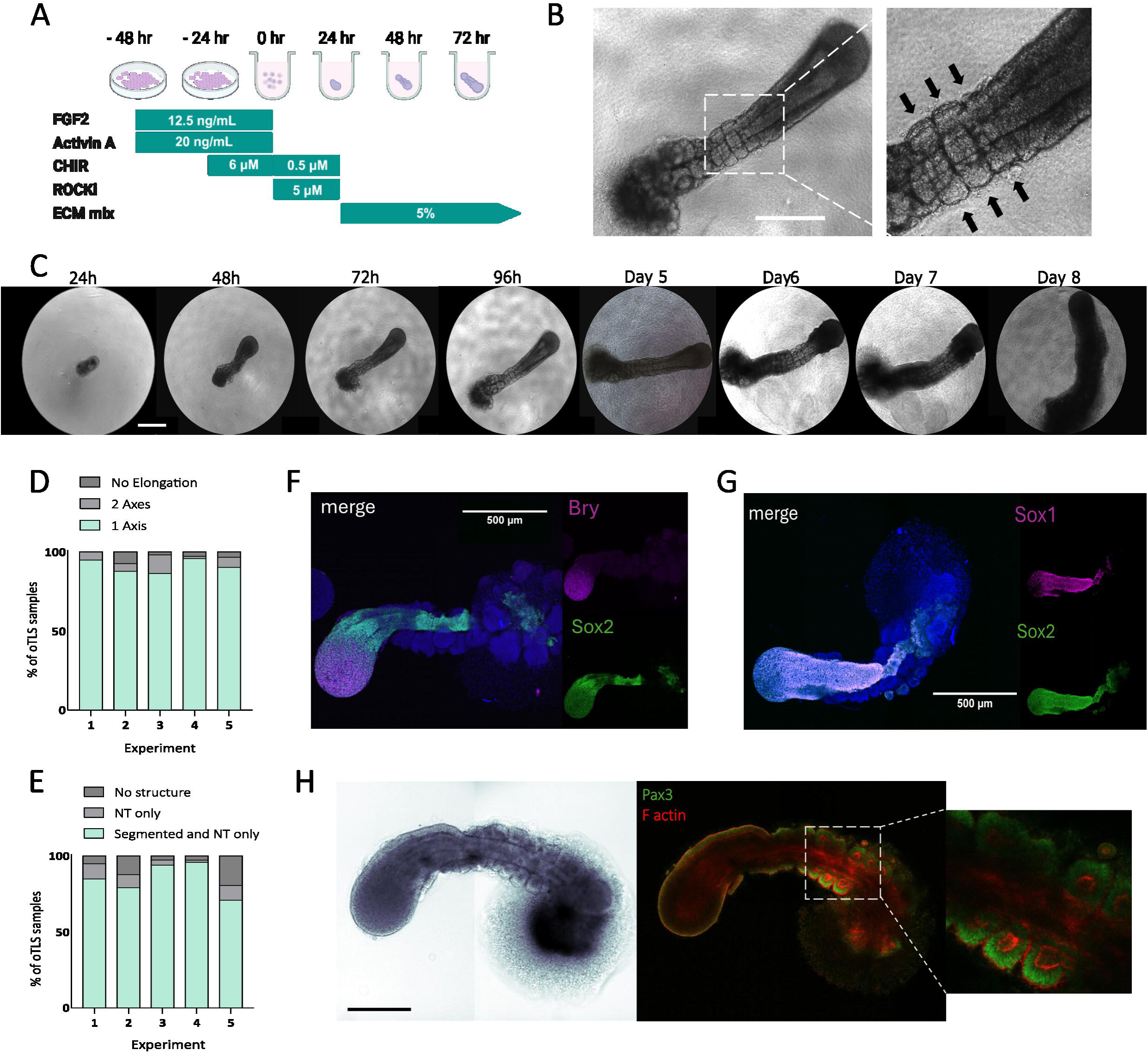
Ovine trunk-like structures robustly develop a neural tube and somites. (A) Graphical representation of established protocol for the formation of ovine TLS. (B) Representative image of a 96h oTLS, showcasing bilateral formation of somite-like structures (indicated by arrows). (C) A representative oTLS imaged every 24h, from day 1 post-aggregation until day 8. (D-E) Quantification of morphogenetic features across experiments: indicating the protocol’s efficiency in terms of (D) axis elongation and (E) neural tube and somite-like morphologies. (F-H) 3D maximum-intensity projection images of immunostained 120□h oTLSs. NMP markers Bry and Sox2 (F) are coexpressed at the posterior end of the oTLS, while Sox2 is also coexpressed with Sox1 (G), corresponding to the NT-like structure, across the midline axis. (H) Confocal section of a 120h oTLS. F-actin staining delineates the apical lumen of somite-like structures, surrounded by Pax3+ immunostained cells. Scale bars 500 µm.

To assess whether the defining cell types of previous TLS models, namely NMPs and their neural tube and somitic derivatives ^9^, are present in the ovine system, we performed an initial characterization of oTLSs by immunofluorescence staining. Bry expression was confined to a discrete domain corresponding to the tailbud region, resembling its localization during axial elongation in vivo. This Bry-positive domain partially overlapped with Sox2 expression, consistent with the presence of an NMP-like population defined by co-expression of Bry and Sox2 ^22^ (Fig. 2F). Bry expression was restricted to the posterior tip and was not detected in a continuous midline axial pattern, as observed in notochord in vivo ^23^. Consistently, Foxa2 expression was not detected, indicating the absence of notochord and definitive endoderm fates in the oTLS model. The central midline of oTLSs exhibited co-expression of Sox2 and Sox1, two canonical markers of early neural identity, indicating a neural tube–like fate (Fig. 2G). Structures flanking the central midline contained a central F-actin–enriched lumen and expressed Pax3 in surrounding cells (Fig. 2H), consistent with dorsal somite and dermomyotome identity. Collectively, the patterned expression of NMP, neural, and somitic markers indicates that oTLSs recapitulate key structural features of early trunk formation in vivo.

### SC RNA-seq characterization of oTLS

To characterize the emergence and maturation of cell types within oTLSs, we performed single-cell RNA sequencing (sc-RNA-seq) across days 2–8 post-aggregation (excluding day 6). Integrated UMAP analysis revealed an increase in cellular diversity as development proceeded, with early time points dominated by progenitor populations and later stages exhibiting a broader spectrum of cell states (Fig. 3A). We identified 14 distinct cell types, with cell-cycle phases evenly distributed across clusters (Fig. S4A). Consistent with the above immunostaining, NMPs and their downstream derivatives, including neural tube–like cells and paraxial mesoderm progressing toward somitic identities, were detected across multiple time points. In addition to these expected lineages, sc-RNA-seq revealed four principal lineage domains: neural, paraxial mesodermal, renal mesodermal, and endothelial lineages. Notably, oTLSs generated a diverse array of neural cell types, including roof plate cells, neural crest populations, glial precursors, midbrain–hindbrain boundary (MHB) progenitors, and postmitotic interneurons (Fig. 3B).

**Figure 3.**
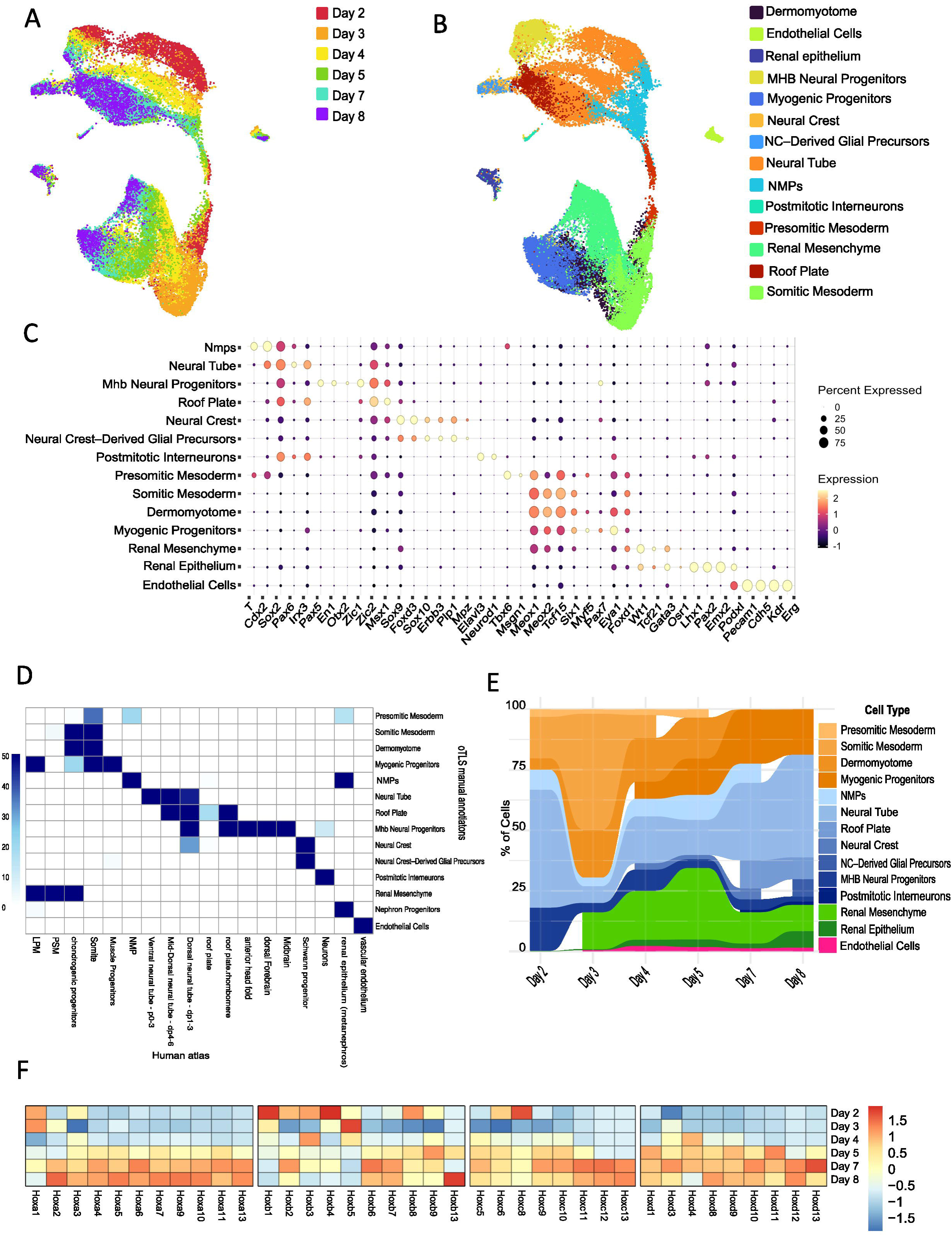
Time-resolved single cell RNA sequencing reveals induction of renal mesoderm, neural crest, MHB progenitors and other advanced cell types in ovine TLS. (A,B) Integrated UMAP of scRNA-seq data from days 2,3,4,5,7 and 8 post aggregation, colored by days (A) or by identified cell types (B). (C) Canonical marker gene expression within identified cell type clusters. (D) Cell type mapping (using SingleR) of ovine TLS cell clusters (columns) to the human atlas dataset (rows). Values signify FDR corrected -log10(Pvalue). (E) Alluvial plot of percentage of the identified cell types over time. (F) Relative expression of HOX genes in oTLS across time points. Expression Z-scores are computed for each gene separately, across the 6 time points.

Across clusters, assigned identities showed strong concordance with canonical marker gene expression profiles (Fig. 3C). To independently assess the accuracy of cell type annotations in the oTLS dataset, we compared our single-cell transcriptomes with an in vivo embryonic reference human atlas ^24^, as well as a mouse extended atlas ^25^ (Fig. 3D, Fig. S4B). This revealed a high degree of concordance between assigned oTLS identities and their expected embryonic counterparts, with only minor deviations observed in a subset of clusters across the dataset.

Lineage composition progressively diversified over time. Neural lineages expanded substantially during oTLS maturation and represented the predominant cell population at later stages. This increase reflects sustained neural differentiation and accumulation of neural derivatives as development progressed (Fig. 3E). Analysis of Hox gene expression across time revealed a progression consistent with embryonic anteroposterior patterning: early Hox paralogs were expressed at earlier stages, whereas progressively more posterior Hox genes were activated at later time points (Fig. 3F), coinciding with observed sustained tailbud elongation.

To assess whether oTLSs recapitulate embryonic stage-dependent metabolic transitions, we quantified the expression of glycolytic and oxidative phosphorylation (OxPhos) gene sets across oTLS developmental time points (Fig. S5A). At early stages, cells expressed both glycolytic and OxPhos-associated genes, consistent with a metabolically bivalent state. As development progressed, glycolytic gene expression increased while OxPhos-associated transcription declined. This shift resembles the in vivo transition where early embryonic tissues rely on bivalent metabolism but progressively adopt glycolysis after implantation ^26^.

We next examined changes in proliferative state over time by quantifying the proportion of cycling cells at each developmental stage (Fig. S5B). At day 2 post-aggregation, more than 90% of cells exhibited gene expression signatures indicative of active proliferation. This proportion decreased steadily during development, dropping below 50% by day 8, coincident with the emergence of differentiated and postmitotic cell populations. These dynamics are consistent with the transition from a progenitor-dominated state toward increasing cell-type maturation observed above.

To investigate whether oTLSs capture vertebrate trunk development dynamics, we analyzed somite formation as a proxy for segmentation-clock activity. Somite rows were visually quantified during days 3–5 post-aggregation, when segmentation was most clearly resolved (Fig. S5C). oTLSs hadan average of 3.7 ± 0.5 somite rows at day 3, which increased to 8.0 ± 0.9 at day 4 and 11.0 ± 0.8 byday 5. Modeling somite accumulation as a linear function of time yielded an estimated segmentation rate of 3.65 somites per day, corresponding to a segmentation periodicity of approximately 6.6 hours per somite pair. Together, this and the above analyses indicate that oTLSs capture multiple dynamic features of early trunk development, including metabolic reprogramming, proliferative decline, and rhythmic segmentation.

### Neuromesodermal progenitor–driven neural and mesodermal differentiation in ovine trunk-like structures

Single-cell transcriptomic and immunofluorescence analyses revealed coordinated neural and mesodermal specification within oTLSs. UMAP visualization identified a discrete population of cells co-expressing the canonical NMP markers Sox2 and Bry positioned at the interface between two major developmental trajectories: a mesodermal lineage comprising paraxial mesoderm and somitic derivatives, and a neural lineage composed of Sox2□Bry□ neural progenitors (Fig. 4A).

**Figure 4.**
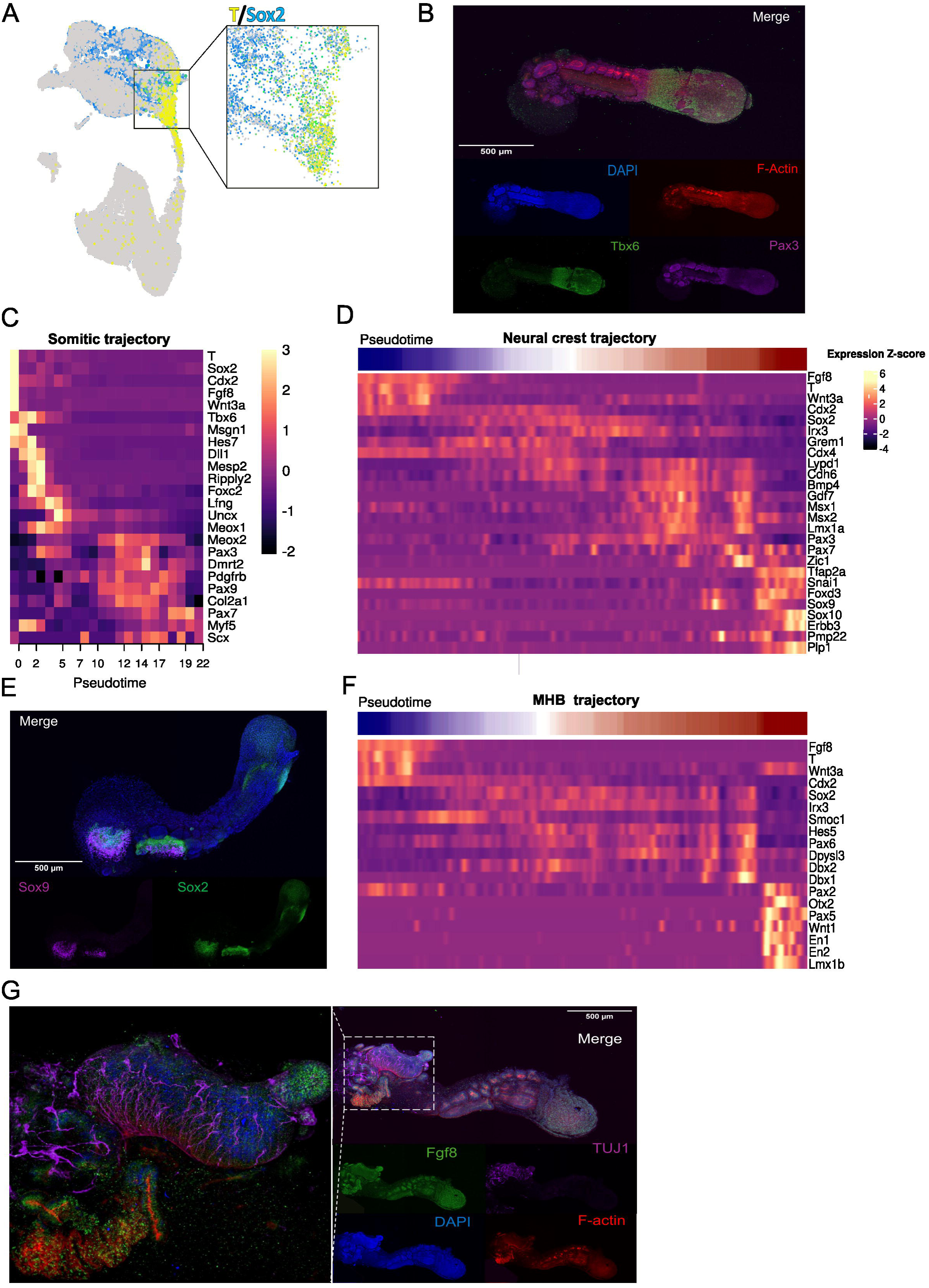
Mesodermal and neural fates in ovine TLS. (A) Integrated UMAP embedding of all single-cell transcriptomes, showing expression of Bry (T) in yellow and Sox2 in blue. Cells co-expressing both markers, representing NMPs, appear as overlapping green points positioned at the interface between neural and mesodermal clusters. (B) 3D immunofluorescence imaging showing Tbx6 expression marking the anterior presomitic mesoderm region of the tailbud, positioned adjacent to Pax3⁺ somite-like domains. F-actin staining highlights epithelialized somite-like units with central lumens. Scale bar 500 µm. (C) Heatmap of stage-specific paraxial mesoderm gene expression in the cell clusters shown in B along pseudotime, demonstrating a gradual and ordered transition from presomitic to somitic and then myogenic identities during oTLS development. (D) Progression of gene marker expression from NMPs to neural tube to neural crest and derived glial precursors along pseudotime. (E) 3D maximum-intensity projection of immunostained day 7 oTLSs. Neural markers Sox9 and Sox2 are expressed adjacently to each other in the anterior region. Scale bar 500 µm. (F) Progression of gene marker expression from NMPs to neural tube to MHB along pseudotime. (G) 3D maximum-intensity projection of immunostained day 7 oTLSs. Secreted protein Fgf8 is expressed throughout the posterior tail, somite-like, and anterior regions, while a condensed anterior domain expresses neuronal marker TUJ1 (magnified, left). Scale bar 500 µm.

Within the paraxial mesoderm, stage-specific marker expression revealed progression from presomitic mesoderm (Msgn1, Tbx6) to somitic (Uncx, Meox2) and myogenic progenitor identities (Myf5, Pax7) (Fig. S7A). At 120h post-aggregation, whole-mount immunofluorescence confirmed spatial organization consistent with early somitogenesis, with Tbx6 expression localized posterior to Pax3□ somite-like domains containing F-actin–enriched lumens (Fig. 4B). Pseudotime analysis supported a continuous transition from presomitic to somitic and myogenic states (Fig. S7B), with sequential marker activation culminating in a myogenic progenitor stage (Fig. 4C).

Within the neural lineage, in addition to axial neural tube populations, differentiated neural subtypes included neural crest cells, glial progenitors, midbrain–hindbrain boundary progenitors, and postmitotic interneurons (Fig. S6C). Pseudotime analysis revealed a structured progression from NMPs to neural tube progenitors, followed by bifurcation toward neural crest and anterior neural identities (Fig. S6D–F), accompanied by sequential activation of corresponding canonical markers (Fig. 4D). Whole-mount immunofluorescence corroborated neural crest identity, with Sox9 detected adjacent to the Sox2□ neuroepithelium in anterior regions of oTLSs (Fig. 4E). Additionally, differentiation toward a midbrain–hindbrain boundary fate was observed across pseudotime (Fig. 4F). Fgf8 expression spanned posterior and anterior regions, while a discrete anterior domain exhibited strong TUJ1 expression (Fig. 4G). TUJ1□ cells formed elongated, aligned extensions arranged in a compact, organized structure, indicative of neuronal differentiation and maturation.

Together, these analyses demonstrate that oTLSs generate NMPs and recapitulate coordinated paraxial mesoderm differentiation and neural lineage diversification, including anterior neural maturation, within a single self-organizing in vitro system

### oTLSs develop mesenchymal and epithelial renal fates

Renal mesodermal populations were first detected on day 3 post-aggregation and progressively diversified during oTLS maturation (Fig. 3A,B). Re-clustering of renal-lineage cells in the single-cell RNA-seq dataset resolved multiple transcriptionally distinct populations expressing canonical renal markers, enabling assignment of nephron-associated and stromal subtypes. These included nephron progenitors (EYA1, FGF10, ITGA8, ALDH1A2), fetal stroma (MEIS1, MEIS2, CRABP1), early nephron epithelium (NCAM1, LHX1, PAX8), podocytes (PTPRO, NPHS1, PODXL), proximal tubules (LRP2, JAG1, FGF8), loop of Henle/distal tubule–like cells (ATP1B1, SLC12A1, PKHD1, KRT18), and collecting duct-like epithelium (GATA3, TFAP2B, EPCAM) (Fig. 5A).

**Figure 5.**
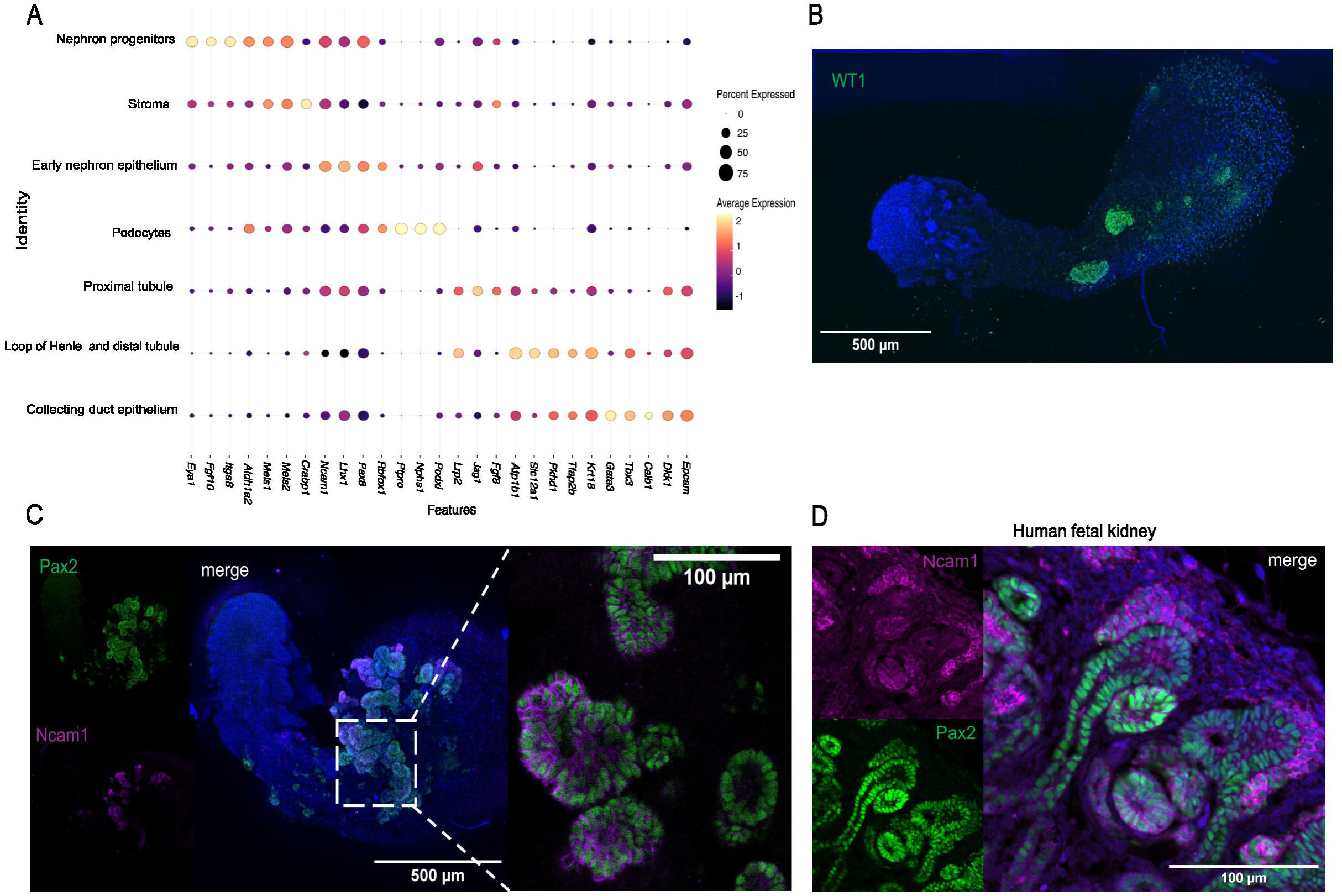
Diversification and spatial organization of renal lineages in oTLSs. (A) Dot plot showing the expression of representative marker genes across the identified renal cell populations. Dot size reflects the proportion of expressing cells and color intensity indicates relative expression level. (B–C) Three-dimensional maximum-intensity projections of immunostained day 7 oTLSs. (B) WT1⁺ domains emerge as bilateral regions in the anterior portion of the oTLS. (C) PAX2 is co-expressed with the membrane marker NCAM1, indicating epithelial differentiation of the metanephric mesenchyme within the anterior region of the oTLS. Right, magnified view of a single Z-slice projection. Scale bars, 500□µm; 100□µm (magnified view). (D) NCAM1⁺ PAX2⁺ immunostaining of week 20 human fetal kidney epithelium. Scale bar 100□µm.

To validate these annotations and assess spatial organization, whole-mount immunostaining for renal markers was performed at day 7 post-aggregation. Bilateral WT1⁺ progenitor populations were observed as lateral domains flanking the medial axis, recapitulating the mediolateral positioning of the kidney field in vivo (Fig. 5B). In the anterior oTLS region, metanephric mesenchyme–like renal progenitors co-expressing nuclear Pax2 and membranous NCAM1 ^27^ were detected (Fig. 5C). These cells formed epithelialized rosette-like structures, consistent with early nephron epithelialization ^28^ and comparable to NCAM1⁺ Pax2⁺ patterning found in human fetal kidney (Fig. 5D). These results indicate that oTLSs generate spatially organized, intermediate mesoderm–derived kidney primordia and recapitulate aspects of early nephrogenic specification and maturation within a trunk-like context.

Together, our findings establish ovine trunk-like structures as a robust, self-organizing in vitro model of post-gastrulation trunk development that integrates axial patterning, neuromesodermal progenitor dynamics, multi-lineage diversification, and early organogenesis within a single in-vitro system.

## Discussion

Mammalian embryogenesis is governed by conserved programs of cell fate specification, morphogenesis, and tissue organization. In recent years, stem cell–based embryo models have enabled unprecedented experimental access to early developmental processes that are otherwise inaccessible in vivo ^29, 4^. However, these systems have been largely restricted to mouse and human species. Here, we extend these models to ungulates, demonstrating the self-organization of ovine and porcine pluripotent stem cells into three-dimensional embryo-like structures recapitulating key aspects of post-gastrulation events. The generation of gastruloids in both species and of trunk-like structures in ovine establish the first in vitro models of ungulate post-gastrulation embryogenesis, enabling comparative analyses of mammalian developmental programs beyond traditional model organisms.

Ovine and porcine gastruloids generated here recapitulate early morphogenetic events, including axial elongation, germ layer specification, and spatial patterning. However, as observed in mouse and human gastruloids, their developmental progression is limited, and they do not autonomously advance to higher-order embryonic structures. Embedding ovine aggregates in a low-percentage ECM environment at defined developmental windows enabled continued morphogenesis and the emergence of trunk-like structures. This transition mirrors findings in mouse and human systems and highlights the importance of the extracellular environment in enabling post-gastrulation tissue organization ^20, 12, 11^.

Transcriptomic analysis showed that ovine gastruloids and early oTLSs generate similar progenitor populations, including neural and paraxial mesoderm derivatives. Their primary differences are found in spatial organization and maturation rather than in progenitor lineage specification. ECM embedding did not significantly alter early fate decisions but instead enabled coordinated morphogenetic progression of progenitor populations into epithelialized neural tube–like structures, segmented somites, and a posterior elongation zone. Together, these results support the notion that ECM signals act predominantly at the tissue level, promoting morphogenesis and structural coherence that support further lineage development rather than introducing new lineages. Consistent with this interpretation, previous studies in mouse TLSs have shown that ECM embedding induces expression of adhesion and cell–ECM interaction pathways, including integrins and cadherins, which are essential for somite epithelialization and neural tube formation in vivo ^20^. The oTLS system therefore reinforces the view that ECM functions not merely as a passive scaffold but as an active regulatory cue that enables embryonic progenitors to execute conserved morphogenetic programs.

The formation of oTLSs demonstrates that ungulate PSCs retain an intrinsic capacity to self-organize into complex, multi-lineage trunk architectures in vitro, even in the absence of extraembryonic tissues. Using time-resolved single-cell transcriptomics, we show that oTLSs generate a broad spectrum of neural, paraxial, intermediate mesodermal, endothelial, and early renal populations arranged along an anteroposterior axis. Notably, oTLSs give rise to dorsal neural derivatives, including neural crest, roof plate, and differentiated neuronal populations, as well as anterior neural identities such as MHB–like domains, which have not been robustly observed in previously described comparable epiblast-only, single-aggregate systems. The emergence of early renal progenitors and spatially organized epithelializing nephric populations further illustrates the capacity of oTLSs to recapitulate aspects of early organogenesis within a unified embryonic context. These findings suggest that oTLSs provide a particularly integrative model for studying cross-germ layer interactions, axial patterning, and early tissue assembly during post-gastrulation development. Nevertheless, the lineage repertoire generated by the models presented here remains constrained, as fates such as endodermal and lateral plate mesoderm derivatives are absent. Primordial germ cells are also absent, as expected given the primed pluripotent state of the ovine and porcine stem cells used in this study ^30^.

A striking feature of the oTLS system is robustness. oTLSs tolerate a wide range of initial aggregate sizes, maintaining axial elongation and segmentation across starting cell numbers spanning more than an order of magnitude. This contrasts with mouse TLSs, which exhibit a narrow size window for successful development and frequently lose axial organization outside this range. In addition, oTLSs remain structurally intact for extended periods, continuing to develop for up to 8–10 days, substantially longer than reported for mouse or human TLS systems. This enhanced robustness may reflect species-specific aspects of ungulate embryogenesis: in vivo, sheep and pigs undergo prolonged pre- and peri-implantation development, with implantation occurring later relative to gastrulation than in mouse or human.

This extended free-floating phase may underlie the sustained morphogenesis we observe under in vitro conditions. While further comparative work will be required to disentangle species-intrinsic properties from culture-dependent effects, these findings highlight the value of extending SEMs to diverse mammalian systems.

Single-cell RNA sequencing revealed marked differences in mesodermal patterning between ovine and porcine gastruloids. Ovine gastruloids formed a defined NMP population that gave rise to neural and paraxial mesoderm lineages, consistent with coordinated trunk elongation. In contrast, porcine gastruloids lacked a clear NMP population and instead exhibited an axial mesoderm–like identity marked by Foxa2 and Shh expression, with Bry localized to an internal elongated domain rather than the posterior pole. This divergence likely underlies the distinct morphogenetic outcomes observed between species and suggests differential responsiveness of ovine and porcine pluripotent stem cells to WNT, FGF, and Nodal signaling under the applied conditions, with porcine cells requiring optimized inputs to support NMP maintenance and trunk morphogenesis.

This study has several limitations. Only a single pluripotent stem cell line per species was analyzed, preventing assessment of inter-line variability. In addition, the lack of a comprehensive ovine embryonic single-cell reference atlas required cross-species annotation using human and mouse datasets, introducing uncertainty due to evolutionary divergence. Furthermore, all experiments were performed under hypoxic conditions, which better approximate early embryonic environments but complicate direct comparison with normoxic mouse and human models. Notably, although hypoxia has been reported to induce ventralization and notochord-like fates in mouse trunk-like structures ^31^, these outcomes were not observed here, indicating that hypoxia-dependent effects are context- and protocol-dependent. Despite these limitations, this work establishes ungulate SEMs as tractable platforms for comparative mammalian developmental biology. By demonstrating that large-mammal PSCs can self-organize into complex trunk-like structures with extended stability and broad lineage representation, this study expands the scope of in vitro embryology beyond traditional model organisms. Future refinements incorporating additional signaling modulation, oxygen control, and expanded reference datasets will further enhance the utility of ungulate SEMs for dissecting conserved and species-specific principles of mammalian development.

## Materials and methods

### Pluripotent stem cell culture

Porcine and ovine embryonic disc–like stem cells (EDSCs) were kindly provided by the Ramiro Alberio laboratory (University of Nottingham, UK) and maintained in a primed pluripotent state under feeder-free and serum-free conditions, as described in ^16^. Briefly, cells were cultured in AFX medium consisting of an N2B27 basal medium, supplemented with Activin A (20 ng mL⁻¹) (120-14E-50, Rhenium), Heat stable FGF2 (12.5 ng mL⁻¹) (100-18BHS-50UG, Rhenium), and the tankyrase inhibitor XAV939 (2 µM) (Sigma X3004, Merck). N2B27 medium was prepared by combining a 1:1 mixture of DMEM/F12 (Gibco 21103049) and Neurobasal medium (Gibco11320074). The basal medium was supplemented with 1× N2 supplement (Gibco17502001), 1× B27 supplement (Gibco17504001), 1x GlutaMAX (Gibco 35050038), 0.05 mM β-mercaptoethanol, and 1% penicillin–streptomycin. The medium was sterile-filtered using a 0.22 µm filter, aliquoted as needed, and stored at 4°C for up to two weeks. Medium was equilibrated to 37°C prior to use. Culture dishes were coated with laminin (10 µg mL⁻¹) (Sigma L2020, Merck) and fibronectin (16.7 µg mL⁻¹) (Sigma F1141, Merck).

Cells were maintained at 38.5 °C in a tri-gas incubator set to 7% CO□ and 5% O□. Cultures were fed daily with freshly supplemented AFX medium. Cells were passaged before reaching full confluence using gentle dissociation with Accutase and replated as small cell clumps to preserve pluripotency and viability. Following dissociation, cells were pelleted by centrifugation and resuspended in fresh medium prior to replating. The ROCK inhibitor Y-27632 was used during thawing and passaging of porcine EDSCs but was not applied to ovine culture passages.

For cryopreservation, cells were dissociated into small clumps and resuspended in NutriFreeze™ D10 cryopreservation medium (05-713-1C, Sartorius) prior to controlled-rate freezing and long-term storage in liquid nitrogen. All cell lines were routinely tested for mycoplasma contamination by PCR and were consistently negative.

### Gastruloid formation

Ovine and porcine gastruloids were generated from pluripotent embryonic disc–like stem cells (EDSCs) maintained in AFX medium. Differentiation was initiated when cultures reached ∼40% confluence, typically one day after passaging. To promote transition to a gastruloid-competent state, XAV939 was withdrawn from the culture medium 48h prior to aggregation, while Activin A and FGF2 were maintained at standard AFX concentrations. Twenty-four hours later, cells were treated with the WNT pathway activator CHIR99021 (SML1046-5MG, Merck). Based on titration experiments, CHIR99021 was used at 6 µM for all subsequent experiments.

After 24h of CHIR99021 pretreatment, cells were dissociated using Accutase (Sigma A6964, Merck), and aggregated in ultra-low-attachment, U-bottom 96-well plates (650970 Greiner Bio-one). Dissociation was quenched with DMEM/F12 supplemented with BSA, and cells were pelleted by centrifugation prior to resuspension in N2B27 medium containing the ROCK inhibitor Y-27632 (5 µM). Cells were counted and seeded at 4,000 cells per well in 50 µL.

Following aggregation, compact spheroids formed within 24h and displayed early symmetry breaking. At this stage, an additional 150 µL of N2B27 medium was added per well, and aggregates were cultured further to allow axial elongation. Gastruloids were collected at indicated time points for downstream analyses.

### Generation of ovine trunk-like structures

Ovine trunk-like structures (oTLSs) were generated from pluripotent embryonic disc–like stem cells (EDSCs) using the same initial pluripotency exit and pre-treatment steps as described for gastruloid formation. Briefly, differentiation was initiated when cultures reached approximately 40% confluence. XAV939 was withdrawn from AFX medium 48h prior to aggregation, while Activin A and FGF2 were maintained at standard concentrations. After 24h, Activin A and FGF2 were refreshed and CHIR99021 was added at 6 µM.

Following 24h of CHIR99021 pretreatment, cells were dissociated, counted, and aggregated in ultra–low-attachment, U-bottom 96-well plates as described for gastruloid formation, with the following modifications. Cells were seeded at 4,000 cells per well in 50 µL of N2B27 medium supplemented with the ROCK inhibitor Y-27632 (5 µM) and CHIR99021 (0.5 µM).

After 24h, compact aggregates had formed and displayed early symmetry breaking. At this stage, 150 µL of N2B27 medium supplemented with 5% (v/v) growth factor–reduced Geltrex was added to each well to support continued morphogenesis and post-gastrulation development. oTLSs were cultured for up to 8 days, during which they underwent sustained axial elongation and adopted trunk-like morphologies, with somite-like segmentation becoming apparent by approximately 72h.

For cultures extending beyond 4 days, half of the medium was replaced with fresh N2B27 supplemented with 5% Geltrex on days 4 and 6 to maintain structural integrity and long-term growth.

### Whole mount Immunostaining

Samples were collected and washed repeatedly in PBS to remove residual extracellular matrix prior to fixation. Fixation was performed in 4% paraformaldehyde at 4 °C for 1 h with gentle agitation. Throughout the staining procedure, samples were handled in U-bottom wells to minimize sample loss during solution exchanges.

After fixation, samples were washed in PBS and permeabilized in PBST (PBS containing 1% Triton X-100) for 1 h at room temperature. Samples were then blocked overnight at 4 °C in PBST supplemented with 5% fetal bovine serum. Primary antibodies were diluted in blocking buffer according to the manufacturers’ recommendations and incubated with samples for 72h at 4 °C with gentle rocking.

Following primary antibody incubation, samples were washed extensively in blocking buffer and PBST and incubated overnight in blocking buffer. Samples were then incubated with fluorophore-conjugated secondary antibodies diluted 1:1000 in blocking buffer for 24h at 4 °C. After washing, nuclei were counterstained with Hoechst (1:1000 in PBS) for 40 min at room temperature. Samples were washed in PBS and stored at 4 °C in PBS until imaging.

Antibodies used in this study:

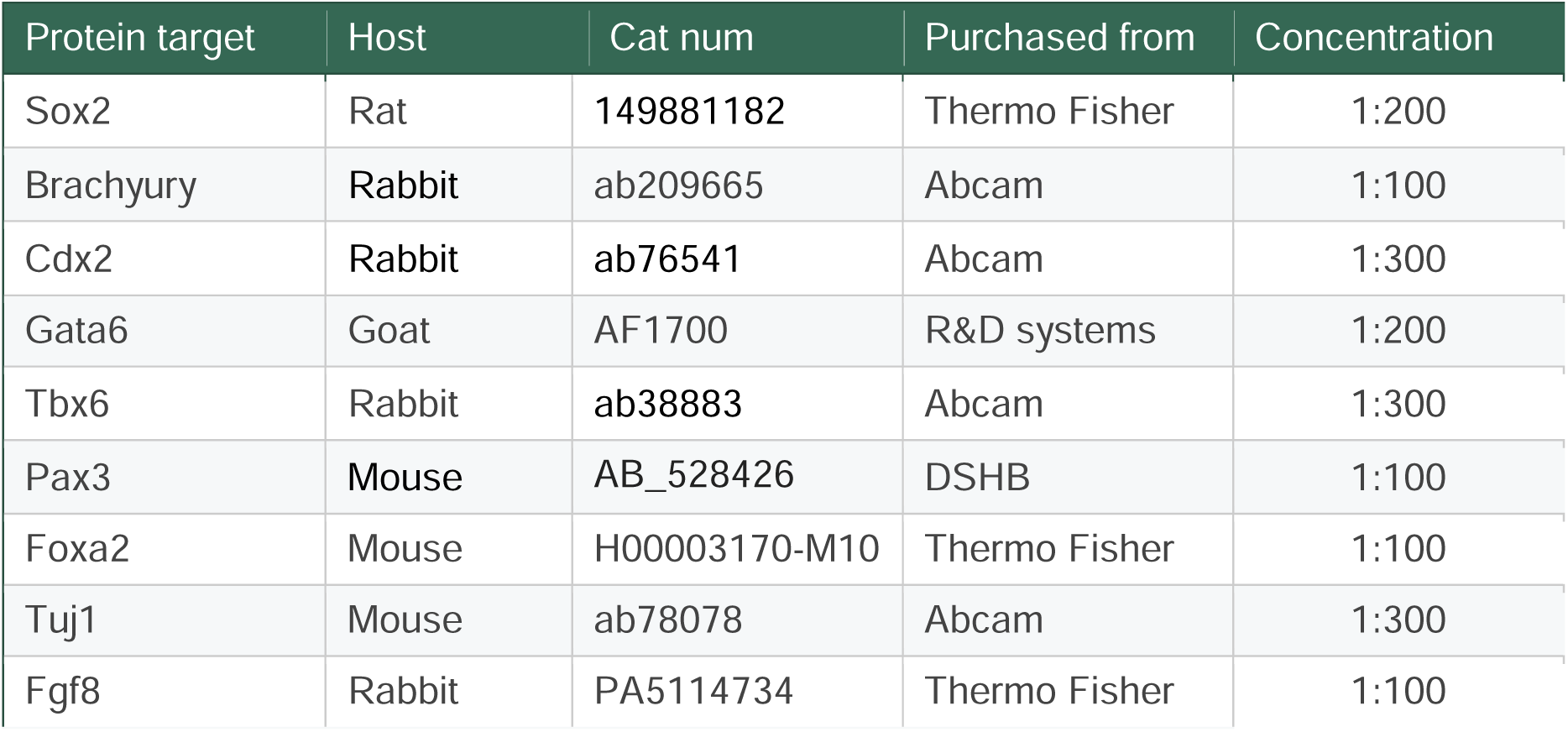
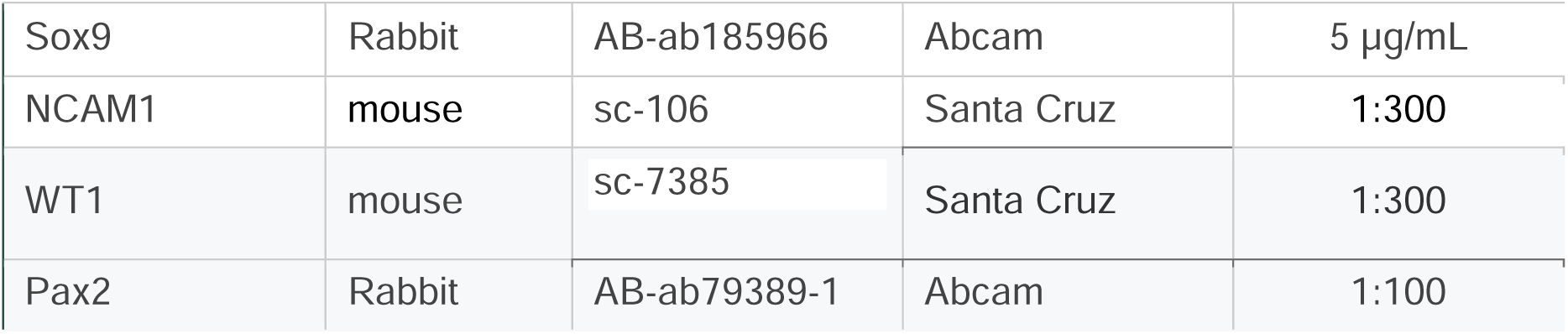

| Protein target | Host | Cat num | Purchased from | Concentration |
| --- | --- | --- | --- | --- |
| Sox2 | Rat | 149881182 | Thermo Fisher | 1:200 |
| Brachyury | Rabbit | ab209665 | Abcam | 1:100 |
| Cdx2 | Rabbit | ab76541 | Abcam | 1:300 |
| Gata6 | Goat | AF1700 | R&D systems | 1:200 |
| Tbx6 | Rabbit | ab38883 | Abcam | 1:300 |
| Pax3 | Mouse | AB_528426 | DSHB | 1:100 |
| Foxa2 | Mouse | H00003170-M10 | Thermo Fisher | 1:100 |
| Tuj1 | Mouse | ab78078 | Abcam | 1:300 |
| Fgf8 | Rabbit | PA5114734 | Thermo Fisher | 1:100 |
| Sox9 | Rabbit | AB-ab185966 | Abcam | 5 µg/mL |
| NCAM1 | mouse | sc-106 | Santa Cruz | 1:300 |
| WT1 | mouse | sc-7385 | Santa Cruz | 1:300 |
| Pax2 | Rabbit | AB-ab79389-1 | Abcam | 1:100 |

### Confocal and epifluorescence microscopy

Imaging was performed using a BC43 spinning disk confocal microscope (Andor, Oxford Instruments). Samples were imaged using 10x and 20x air objectives, depending on the required field of view and resolution. Z-stacks were acquired at 5–7 µm intervals across the full sample depth to capture three-dimensional tissue organization. Excitation was provided by solid-state lasers at 405, 488, 561, and 640 nm, and fluorescence emission was collected using standard filter sets corresponding to DAPI, Alexa Fluor 488, Alexa Fluor 568, and Alexa Fluor 647.

Samples were mounted individually in a small volume of PBS on glass-bottom dishes for imaging. For two-dimensional stained cells adhered to plastic culture wells, images were acquired using epifluorescence settings.

Post-acquisition image processing, including maximum intensity projection, background subtraction, and linear brightness and contrast adjustments, was performed uniformly using Andor Fusion software or FIJI–ImageJ (NIH). Scale bars, annotations, and final figure preparation were carried out in FIJI–ImageJ.

### Brightfield imaging and morphometric analysis

Live-sample brightfield imaging was performed using an Echo Rebel benchtop microscope (Echo, San Diego, CA, USA). Samples were imaged at room temperature under brightfield illumination using 10x or 20x objectives with the instrument’s integrated digital camera. Images were acquired using identical acquisition settings across conditions within each experiment. Scale bars were generated using the microscope’s built-in calibration system and are shown on all images.

Morphometric measurements, including sample area, length, and width, were obtained using the Rebel Echo image acquisition software. Gastruloid and trunk-like structure outlines and principal axes were manually traced, and quantitative measurements were extracted from these tracings for downstream analysis.

### Human fetal kidney samples and immunofluorescence

Week 20 human fetal kidney samples were obtained from elective pregnancy terminations following written informed consent for tissue collection and subsequent analysis. The study was conducted in accordance with the principles of the Declaration of Helsinki. Ethical approval was granted by the Institutional Review Boards of Shamir Medical Center (approval no. 0132-18-ASF).

Excised tissues were rinsed thoroughly in phosphate-buffered saline (PBS), fixed, embedded in paraffin, and sectioned at 5 μm thickness. Paraffin sections were subjected to antigen retrieval using OmniPrep solution (pH 9.0; Zytomed Systems) at 95□°C for 1□h according to the manufacturer’s instructions. Sections were blocked with Cas-Block solution (Invitrogen ImmunoDetection, 00-8120) for 20□min at room temperature.

Slides were incubated with primary antibodies (listed above) followed by species-appropriate secondary antibodies conjugated to Alexa Fluor dyes (Alexa Fluor 488 anti-rabbit, Alexa Fluor 555 anti-mouse, and Alexa Fluor 647 anti-goat; 1:1200 dilution) for 60□min at room temperature. Nuclei were counterstained using DAPI-containing mounting medium (DAPI Fluoromount-G; SouthernBiotech, 0100-20).

Fluorescence images were acquired using an Olympus IX83 microscope equipped with an Olympus DP80 camera. Image processing was performed using Fiji.

### Single-cell RNA sequencing and analysis

#### Single-cell dissociation and library preparation

Gastruloids and trunk-like structures were generated as described above. For single-cell RNA sequencing (scRNA-seq), pools of 48 porcine gastruloids (days 2 and 3), 48 ovine gastruloids (days 2 and 3), and 32 ovine trunk-like structures (days 2, 3, 4, 5, 7, and 8) were collected per time point. Structures were transferred individually using cut pipette tips, washed repeatedly in ice-cold PBS to remove extracellular matrix residues, and enzymatically dissociated using TrypLE Express at 37 °C with intermittent gentle trituration until a single-cell suspension was obtained. Enzymatic reactions were quenched with PBS containing BSA, and cells were washed prior to pooling by time point.

Cells were pelleted by centrifugation, resuspended in PBS containing 0.4% BSA, and viable cells were counted. Single-cell suspensions were immediately processed using the Chromium GEM-X Universal 3′ Gene Expression v4 platform (10x Genomics) according to the manufacturer’s instructions, including on-chip sample multiplexing. Libraries were prepared using the GEM-X Universal 3′ Gene Expression v4 4-plex kit and sequenced on an Illumina NextSeq 2000 system.

#### Preprocessing and alignment

Base call files were converted to FASTQ format using bcl2fastq (v2.20.0). Reads were aligned using Cell Ranger (v9.0.1). A sheep reference genome was generated using Cell Ranger mkref based on the ARS-UI_Ramb_v3 assembly and corresponding gene annotation. Filtered feature–barcode matrices were generated using Cell Ranger multi and used for downstream analyses.

#### Quality control and data processing

Single-cell datasets were analyzed using Seurat (v5.3.0). Cells with low gene counts, high mitochondrial or ribosomal transcript content, or extreme UMI counts were excluded based on per-sample quality metric distributions. Putative doublets were identified and removed using DoubletFinder (v2.0.6). Each dataset was initially processed independently and then merged by species and developmental stage. Batch correction was applied to porcine gastruloid datasets using anchor-based integration.

Merged datasets were log-normalized, variable genes were identified using the “vst” method, and data were scaled while regressing out cell cycle phase, UMI counts, and ribosomal gene expression. Principal component analysis was performed, and the first 20 principal components were used for nearest-neighbor graph construction, clustering, and UMAP dimensionality reduction.

#### Cell type annotation

Cell types were annotated based on differential gene expression and canonical marker expression. Differentially expressed genes were identified for each cluster using a Wilcoxon rank-sum test with multiple testing correction. Cluster identities were assigned based on known markers of germ layers, tissues, and progenitor populations, and validated by assessing co-expression of additional canonical markers. Only clusters supported by consistent marker expression and developmental plausibility were assigned specific identities; ambiguous clusters were labeled as unclassified or assigned broader lineage identities.

#### Cross-species annotation using SingleR

To support and cross-validate manual annotations, SingleR (v2.8.0) was applied using a mouse^25^ embryonic reference atlas spanning embryonic days E6.5–E9.5, and a human embryonic reference atlas spanning Carnegie stages 12-16 (4-6 weeks) ^24^. Ovine and Human gene symbols were mapped to mouse orthologs prior to analysis. SingleR was run on both raw and batch-corrected expression matrices. Cells with low annotation confidence were left unlabeled. SingleR assignments were interpreted at the lineage level and used as supportive evidence rather than definitive labels.

#### Trajectory and pseudotime analysis

Lineage-specific developmental trajectories were inferred using Monocle 3. Cells were subset by lineage prior to analysis. For neural lineages, only later time points (days 5, 7, and 8) were included to minimize batch-driven discontinuities. The Myogenic lineage was analyzed using all relevant time points.

Lineage-specific Seurat objects were reprocessed and converted to Monocle cell_data_set objects. Root states were defined based on biological context, corresponding to neuromesodermal progenitors for neural and myogenic lineages. Pseudotime values were assigned to individual cells.

For the myogenic lineage, pseudotime values were smoothed across nearest neighbors and binned to visualize continuous transcriptional dynamics. The Neural trajectories were analyzed without smoothing. In the neural lineage, bifurcating trajectories were identified downstream of neuromesodermal progenitors, corresponding to dorsal neural crest–associated and anterior neural branches. Branch-specific pseudotime analyses were performed to examine transcriptional dynamics and axial patterning, including HOX gene expression.

### Quantification and statistical analysis

Statistical analyses were performed using GraphPad Prism (v8.0.1, GraphPad Software). Data are presented as mean ± standard deviation unless otherwise indicated. Comparisons between multiple groups were performed using two-way analysis of variance (ANOVA) followed by Tukey’s post hoc test for multiple comparisons. A p value < 0.05 was considered statistically significant. The number of independent biological replicates is indicated in the figure legends.

### Graphics

Protocol graphical representation and additional schemes were created with BioRender.com.

## Acknowledgements

We thank J. Veenvliet, O. Revah, D. Sela-Donenfeld, R. Rak, E. Tzahor, and M. Danan-Gotthold for helpful discussions, and R. Alberio and D. Klisch for their kindness and generosity in sharing their cell lines and knowledge. We thank J. Kippen and S. Greggs for rigorous proofreading and editing. This study was supported by the Israel Science Foundation (1491/22) and the Good Food Institute research program.

## Author Contributions

M.H. and I.N. conceived the study. M.H. performed all experiments. M.N. and B.D. contributed fetal human kidney immunostaining and imaging. P.B. wrote all the single-cell RNA seq analysis code. M.H. and P.B. analyzed the data with contributions from M.N. M.H. and I.N. wrote the manuscript with contributions from all authors. I.N. and S.S. oversaw the project.

---

**Supplementary Figure 1. Extent of elongation in ovine gastruloids depends on the strength of WNT pathway activation.** (A) Representative 72h ovine gastruloids under four varying concentrations of CHIR pretreatments. Scale bar 530 µm. (B) Quantification of length-to-width ratio of ovine gastruloids pre-treated with the indicated concentrations of CHIR, over time. (n= 24, **P<0.005).

**Supplementary Figure 2. Establishment and optimization of an ovine trunk-like structure protocol.** (A) Efficiency of elongation and axial tube formation at 72h in samples where geltrex was introduced at 24h vs at 48h. (B) Representative images displaying the difference in morphology at 72h, without the timely addition of 5% ECM matrix and with its addition. Scale bar 200 µm. (C) The effects of a 24-hr break in Wnt inhibition before activation by CHIR and of adding retinoic acid (RA) between 0h-24h on somite and tube formation success. (D) Extent of asphericity at 24h post aggregation predicts the eventual success of oTLS morphology formation at 72h. ***p<0.001. (E) Left - Representative brightfield images of samples with and without ECM matrix addition, under four different concentrations of CHIR pretreatment. Right - magnification of a sample with 6 µM CHIR pretreatment at 72h, showcasing bilateral formation of somite-like structures (pointed by arrows).

**Supplementary Figure 3. oTLS form somites and a neural tube across a wide range of initial cell numbers.** (A) Representative images of 96h oTLSs aggregated from 400, 1000, 2000, 4000 and 8000 cells, all showcasing the formation of tube and flanking somite-like structures and quantification of oTLS log2 of area (B) and length (C) over time across initial cell numbers.

**Supplementary Figure 4.** (A) oTLS integrated UMAP, colored by inferred cell-cycle phase (G1, S, G2/M), demonstrating homogeneous distribution across clusters and absence of cell-cycle–driven segregation. (B) Cell type mapping (using SingleR) of ovine TLS cell clusters (columns) to the extended mouse atlas dataset (rows) ^25^. Values signify FDR corrected -log10(Pvalue).

**Supplementary Figure 5. Temporal dynamics of oTLS.** (A) UMAP showing scaled, whole-dataset expression of glycolytic genes (blue) and oxidative phosphorylation genes (yellow) across experimental time points, separated by day post-aggregation. A shift toward increased glycolytic gene expression is observed with progression from early (green) to later (blue) stages. (B) Proportion of proliferative cells over time, showing a steady decline in dividing populations as differentiation proceeds. (C) Quantification of somite row formation from 72□h to 120□h across five oTLSs, used to calculate segmentation clock oscillation periodicity.

**Supplementary Figure 6. Paraxial mesoderm and neural trajectories in oTLS.** (A) UMAP visualization showing expression of canonical markers spanning successive stages of paraxial mesoderm development. Early presomitic mesoderm markers (Msgn1, Tbx6) are followed by somitic markers (Uncx, Meox2) and later myogenic progenitor markers (Myf5, Pax7). (B) Monocle pseudotime trajectory constructed from paraxial mesoderm and myogenic clusters. (C) UMAP visualization shows expression of neural fate marker genes. Otx2 and Lhx2 mark MHB, Sox10 and Foxd3 mark neural crest, Elavl3 and Isl1 mark postmitotic interneurons. (D) Monocle-based pseudotime analysis plot of neural clusters, split for downstream analysis into trajectories leading to the neural crest fates (E) and MHB fates (F).

## References

1. Bazer, F.W., Spencer, T.E., Johnson, G.A., Burghardt, R.C., and Wu, G. (2009). Comparative aspects of implantation. Reproduction 138, 195–209. 10.1530/REP-09-0158.

2. Roberts, R.M., Green, J.A., and Schulz, L.C. (2016). The evolution of the placenta. Reproduction 152, R179–89. 10.1530/REP-16-0325.

3. Simpson, L., and Alberio, R. (2023). Interspecies control of development during mammalian gastrulation. Emerg. Top. Life Sci. 7, 397–408. 10.1042/ETLS20230083.

4. Veenvliet, J.V., and Herrmann, B.G. (2021). Modeling mammalian trunk development in a dish. Dev. Biol. 474, 5–15. 10.1016/j.ydbio.2020.12.015.

5. van den Brink, S.C., Baillie-Johnson, P., Balayo, T., Hadjantonakis, A.-K., Nowotschin, S., Turner, D.A., and Martinez Arias, A. (2014). Symmetry breaking, germ layer specification and axial organisation in aggregates of mouse embryonic stem cells. Development 141, 4231–4242. 10.1242/dev.113001.

6. Moris, N., Anlas, K., van den Brink, S.C., Alemany, A., Schröder, J., Ghimire, S., Balayo, T., van Oudenaarden, A., and Martinez Arias, A. (2020). An in vitro model of early anteroposterior organization during human development. Nature 582, 410–415. 10.1038/s41586-020-2383-9.

7. van den Brink, S.C., and van Oudenaarden, A. (2021). 3D gastruloids: a novel frontier in stem cell-based in vitro modeling of mammalian gastrulation. Trends Cell Biol. 31, 747–759. 10.1016/j.tcb.2021.06.007.

8. Turner, D.A., and Martinez Arias, A. (2024). Three-dimensional stem cell models of mammalian gastrulation. Bioessays 46, e2400123. 10.1002/bies.202400123.

9. Veenvliet, J.V., Bolondi, A., Kretzmer, H., Haut, L., Scholze-Wittler, M., Schifferl, D., Koch, F., Guignard, L., Kumar, A.S., Pustet, M., et al. (2020). Mouse embryonic stem cells self-organize into trunk-like structures with neural tube and somites. Science 370. 10.1126/science.aba4937.

10. van den Brink, S.C., Alemany, A., van Batenburg, V., Moris, N., Blotenburg, M., Vivié, J., Baillie-Johnson, P., Nichols, J., Sonnen, K.F., Martinez Arias, A., et al. (2020). Single-cell and spatial transcriptomics reveal somitogenesis in gastruloids. Nature 582, 405–409. 10.1038/s41586-020-2024-3.

11. Makwana, K., Tilley, L., Chakravarty, P., Thompson, J., Baillie-Benson, P., Rodriguez-Polo, I., and Moris, N. (2025). Modelling co-development between the somites and neural tube in human trunk-like structures. Nat. Cell Biol. 27, 2049–2062. 10.1038/s41556-025-01813-8.

12. Hamazaki, N., Yang, W., Kubo, C.A., Qiu, C., Martin, B.K., Garge, R.K., Regalado, S.G., Nichols, E.K., Pendyala, S., Bradley, N., et al. (2024). Retinoic acid induces human gastruloids with posterior embryo-like structures. Nat. Cell Biol. 26, 1790–1803. 10.1038/s41556-024-01487-8.

13. Balaskas, A., Kraus, I., Özgüldez, H.Ö., Omgba, P.A., Buschow, R., Bolondi, A., Berlad, I., Hanna, J.H., Kretzmer, H., and Bulut-Karslioğlu, A. (2025). An advanced head-to-tail mouse embryo model with hypoxia-mediated neural patterning. BioRxiv. 10.1101/2025.06.17.660116.

14. Navarro, M., Laiz-Quiroga, L., Blüguermann, C., and Mutto, A. (2024). Livestock embryonic stem cells for reproductive biotechniques and genetic improvement. Anim. Reprod. 21, e20240029. 10.1590/1984-3143-AR2024-0029.

15. Xiang, J., Wang, H., Shi, B., Li, J., Liu, D., Wang, K., Wang, Z., Min, Q., Zhao, C., and Pei, D. (2024). Pig blastocyst-like structure models from embryonic stem cells. Cell Discov. 10, 72. 10.1038/s41421-024-00693-w.

16. Kinoshita, M., Kobayashi, T., Planells, B., Klisch, D., Spindlow, D., Masaki, H., Bornelöv, S., Stirparo, G.G., Matsunari, H., Uchikura, A., et al. (2021). Pluripotent stem cells related to embryonic disc exhibit common self-renewal requirements in diverse livestock species. Development 148. 10.1242/dev.199901.

17. Beccari, L., Moris, N., Girgin, M., Turner, D.A., Baillie-Johnson, P., Cossy, A.-C., Lutolf, M.P., Duboule, D., and Arias, A.M. (2018). Multi-axial self-organization properties of mouse embryonic stem cells into gastruloids. Nature 562, 272–276. 10.1038/s41586-018-0578-0.

18. Rossant, J., and Tam, P.P.L. (2017). New Insights into Early Human Development: Lessons for Stem Cell Derivation and Differentiation. Cell Stem Cell 20, 18–28. 10.1016/j.stem.2016.12.004.

19. José-Edwards, D.S., Oda-Ishii, I., Kugler, J.E., Passamaneck, Y.J., Katikala, L., Nibu, Y., and Di Gregorio, A. (2015). Brachyury, Foxa2 and the cis-Regulatory Origins of the Notochord. PLoS Genet. 11, e1005730. 10.1371/journal.pgen.1005730.

20. Veenvliet, J.V., Bolondi, A., Kretzmer, H., Haut, L., Scholze-Wittler, M., Schifferl, D., Koch, F., Guignard, L., Kumar, A.S., Pustet, M., et al. (2020). Mouse embryonic stem cells self-organize into trunk-like structures with neural tube and somites. Science 370. 10.1126/science.aba4937.

21. Farag, N., Sacharen, C., Avni, L., and Nachman, I. (2024). Coordination between endoderm progression and mouse gastruloid elongation controls endodermal morphotype choice. Dev. Cell 59, 2364–2374.e4. 10.1016/j.devcel.2024.05.017.

22. Unable to find information for 763308.

23. Kemmler, C.L., Smolikova, J., Moran, H.R., Mannion, B.J., Knapp, D., Lim, F., Czarkwiani, A., Hermosilla Aguayo, V., Rapp, V., Fitch, O.E., et al. (2023). Conserved enhancers control notochord expression of vertebrate Brachyury. Nat. Commun. 14, 6594. 10.1038/s41467-023-42151-3.

24. Xu, Y., Zhang, T., Zhou, Q., Hu, M., Qi, Y., Xue, Y., Nie, Y., Wang, L., Bao, Z., and Shi, W. (2023). A single-cell transcriptome atlas profiles early organogenesis in human embryos. Nat. Cell Biol. 25, 604–615. 10.1038/s41556-023-01108-w.

25. Imaz-Rosshandler, I., Rode, C., Guibentif, C., Harland, L.T.G., Ton, M.-L.N., Dhapola, P., Keitley, D., Argelaguet, R., Calero-Nieto, F.J., Nichols, J., et al. (2024). Tracking early mammalian organogenesis - prediction and validation of differentiation trajectories at whole organism scale. Development 151. 10.1242/dev.201867.

26. Malkowska, A., Penfold, C., Bergmann, S., and Boroviak, T.E. (2022). A hexa-species transcriptome atlas of mammalian embryogenesis delineates metabolic regulation across three different implantation modes. Nat. Commun. 13, 3407. 10.1038/s41467-022-30194-x.

27. Namestnikov, M., Cohen-Zontag, O., Omer, D., Gnatek, Y., Goldberg, S., Vincent, T., Singh, S., Shiber, Y., Rafaeli Yehudai, T., Volkov, H., et al. (2025). Human fetal kidney organoids model early human nephrogenesis and Notch-driven cell fate. EMBO J. 44, 4681–4719. 10.1038/s44318-025-00504-2.

28. Vincent, T., Nademi, S., Namestnikov, M., Cohen-Zontag, O., Dekel, B., and Freedman, B.S. (2026). Co-induction of stromal and epithelial progenitors for renal regeneration. The Innovation, 101281. 10.1016/j.xinn.2026.101281.

29. Sozen, B., Conkar, D., and Veenvliet, J.V. (2022). Carnegie in 4D? Stem-cell-based models of human embryo development. Semin. Cell Dev. Biol. 131, 44–57. 10.1016/j.semcdb.2022.05.023.

30. Neupane, J., Lubatti, G., Gross-Thebing, T., Ruiz Tejada Segura, M.L., Butler, R., Gross-Thebing, S., Dietmann, S., Scialdone, A., and Surani, M.A. (2025). The emergence of human primordial germ cell-like cells in stem cell-derived gastruloids. Sci. Adv. 11, eado1350. 10.1126/sciadv.ado1350.

31. López-Anguita, N., Gassaloglu, S.I., Stötzel, M., Bolondi, A., Conkar, D., Typou, M., Buschow, R., Veenvliet, J.V., and Bulut-Karslioglu, A. (2022). Hypoxia induces an early primitive streak signature, enhancing spontaneous elongation and lineage representation in gastruloids. Development 149. 10.1242/dev.200679.

