## Supplementary figures and images for "Robust self-organization of livestock pluripotent stem cells into post-gastrulation embryo models with advanced neuronal and mesodermal structures"

### Supplementary Figure 1

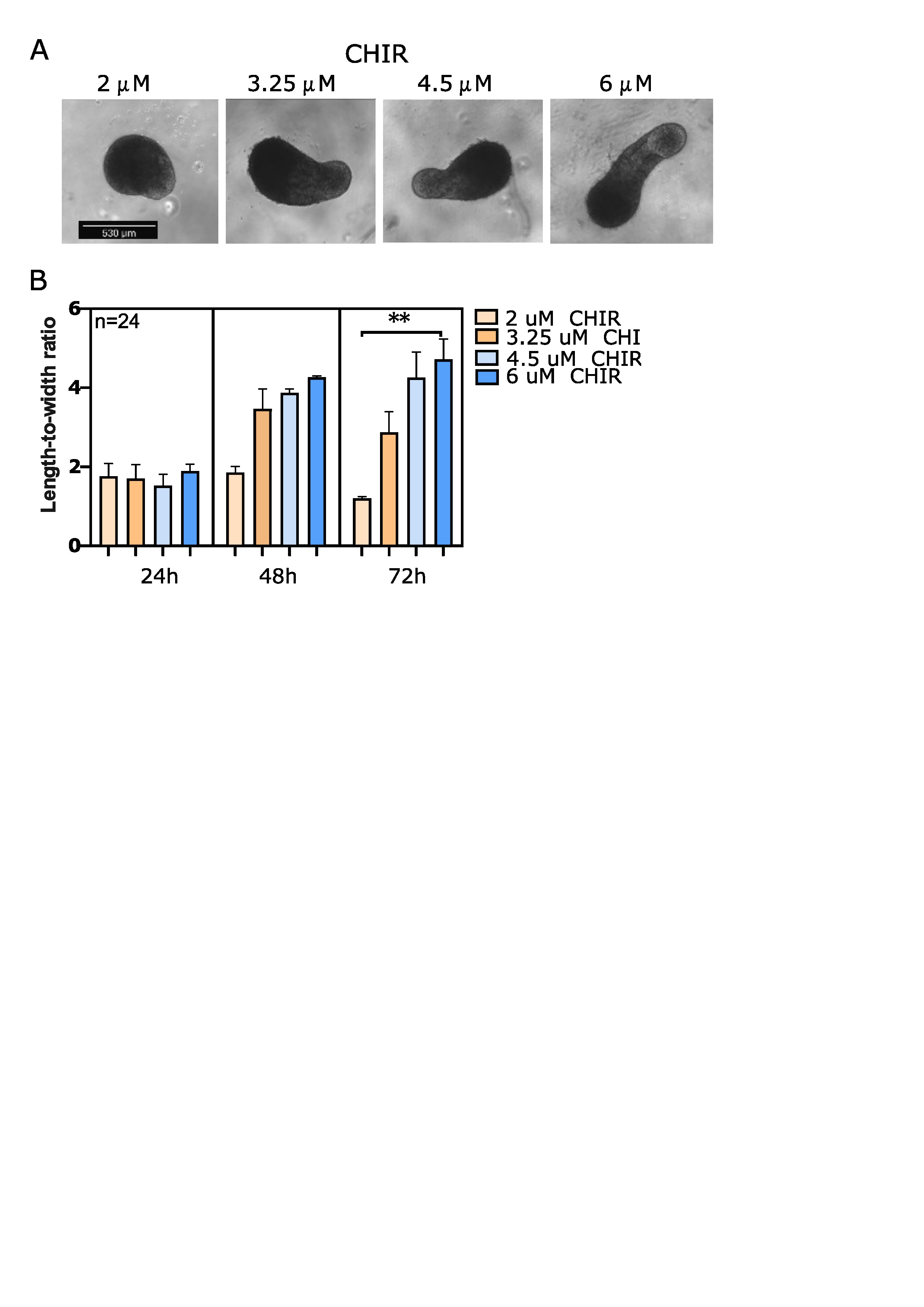

### Supplementary Figure 2

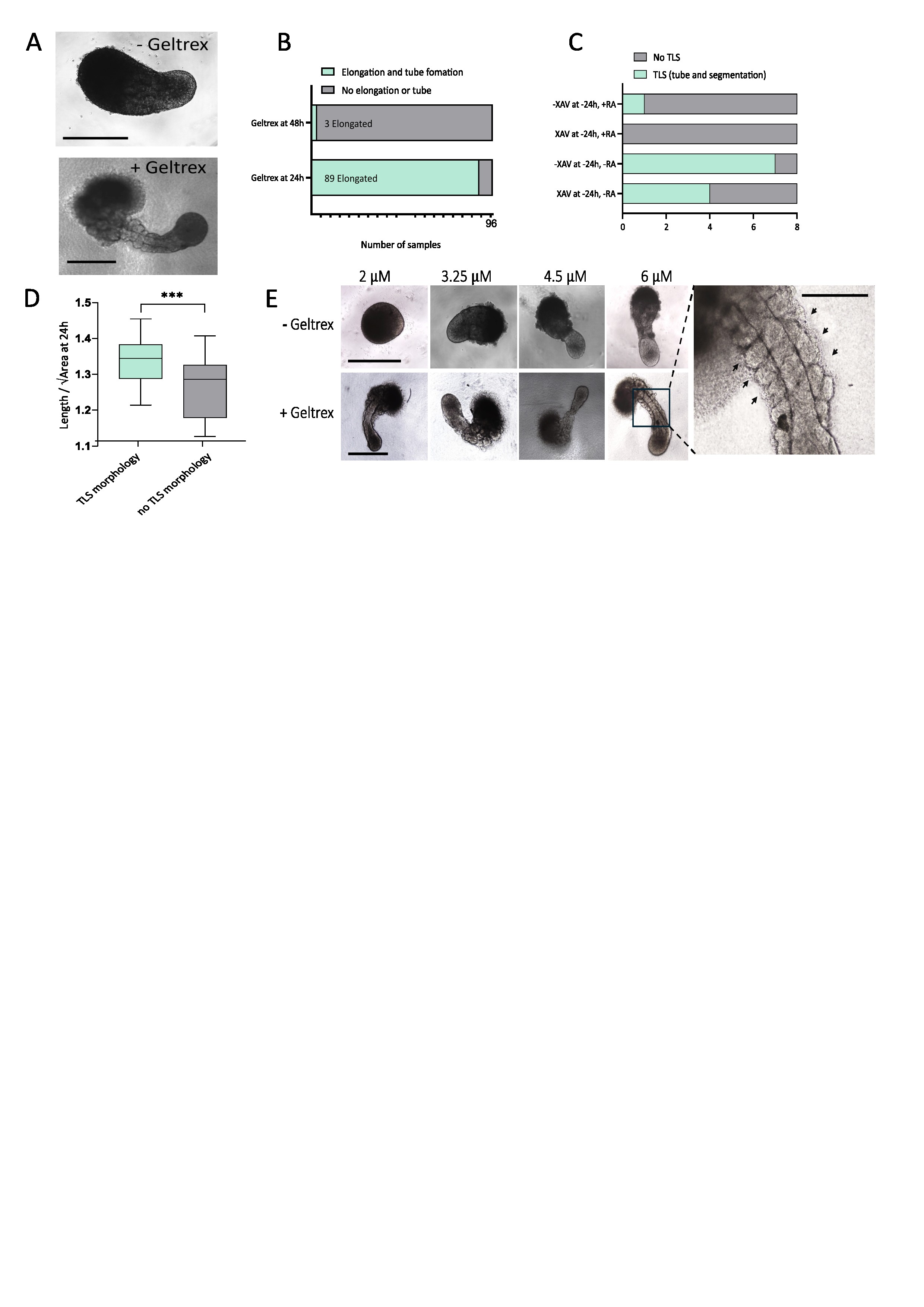

### Supplementary Figure 3

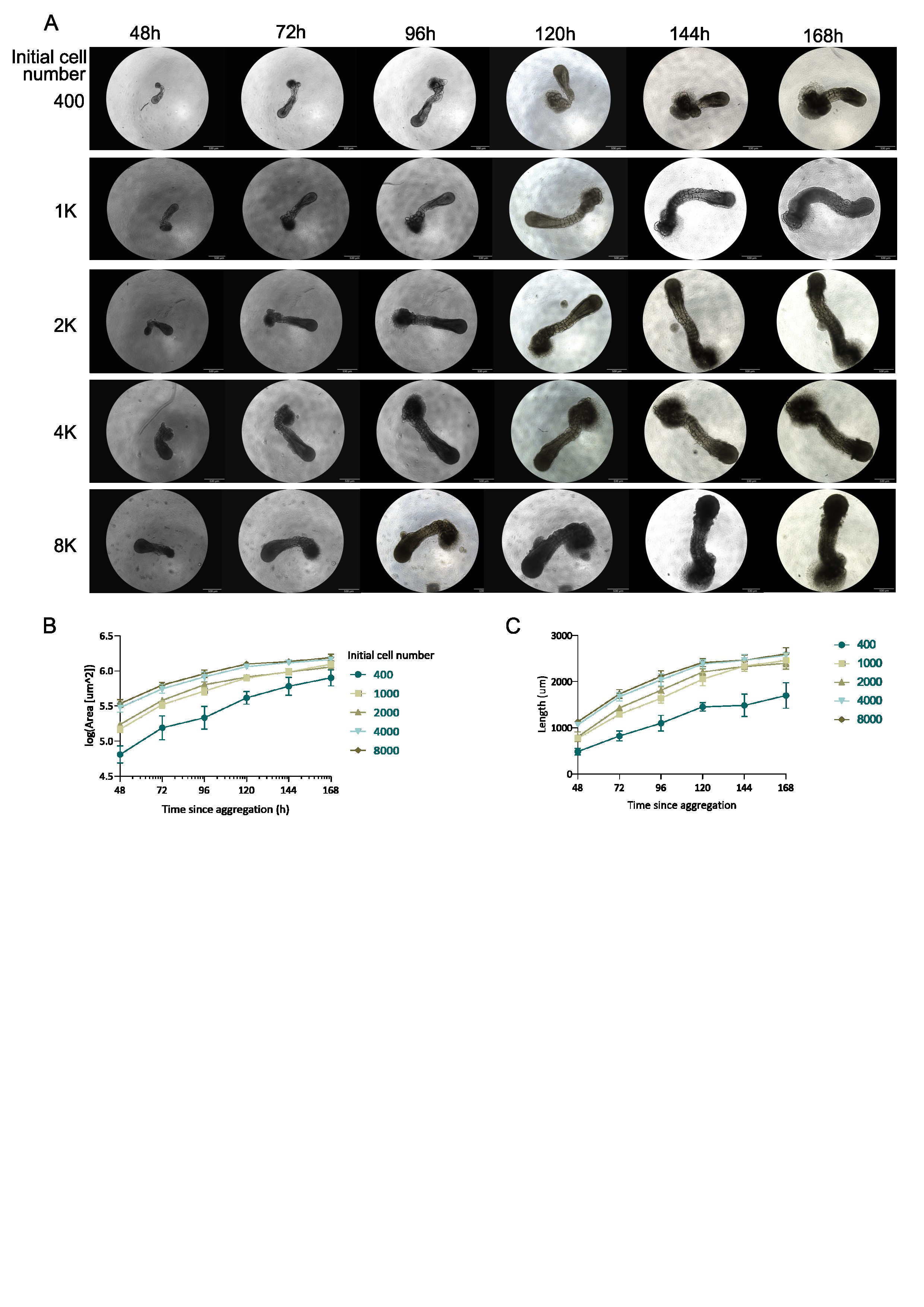

### Supplementary Figure 4

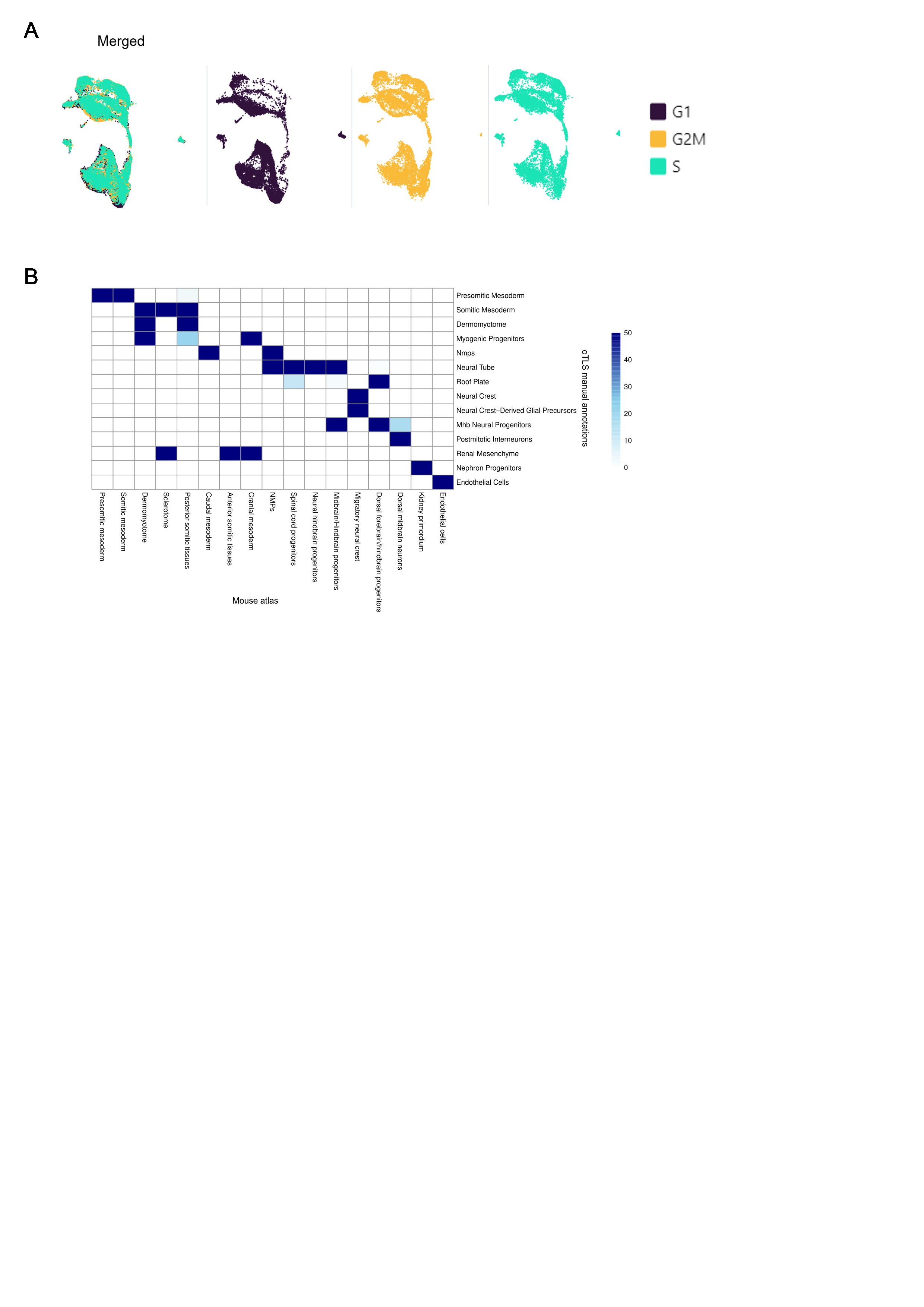

### Supplementary Figure 5

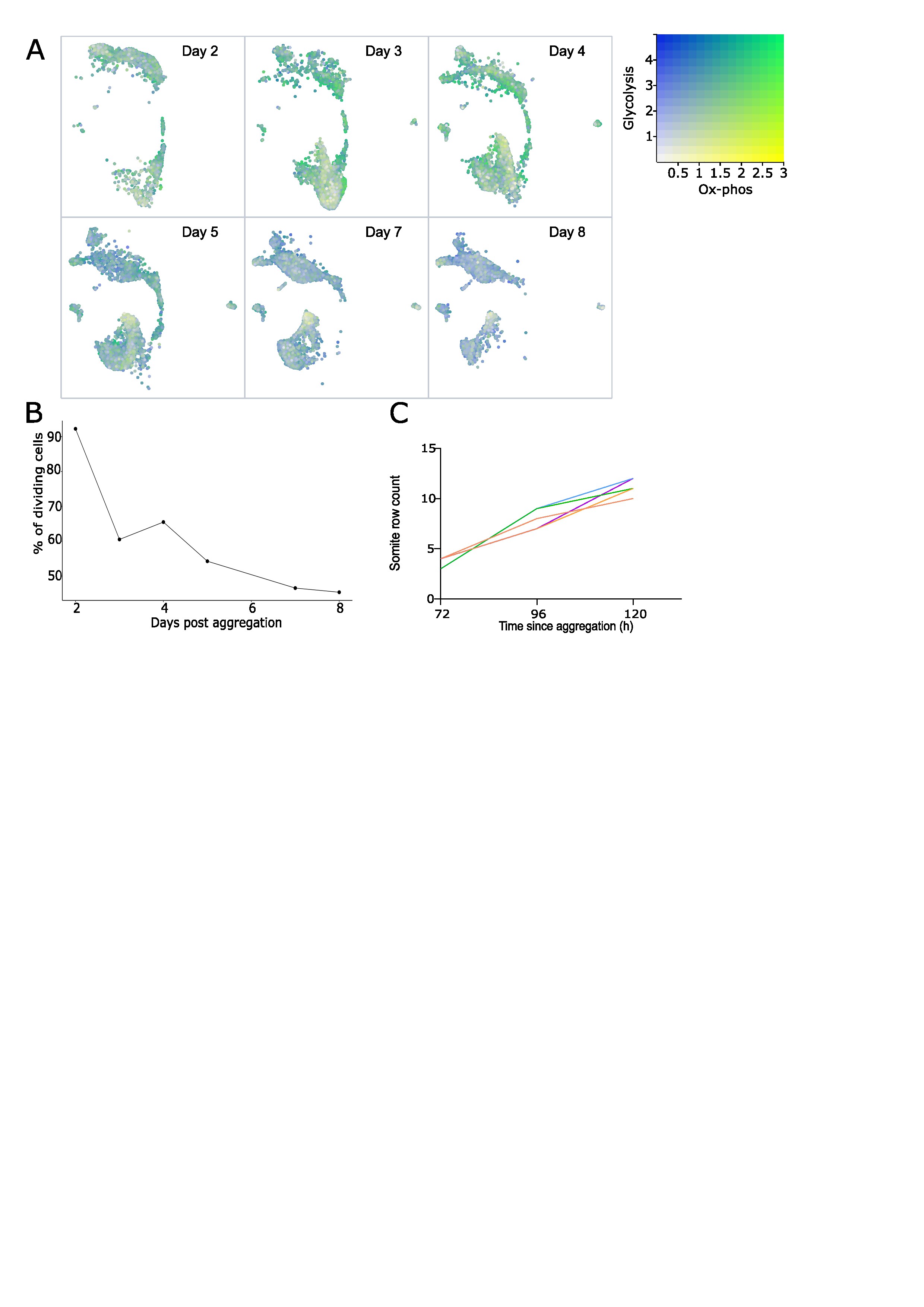

### Supplementary Figure 6

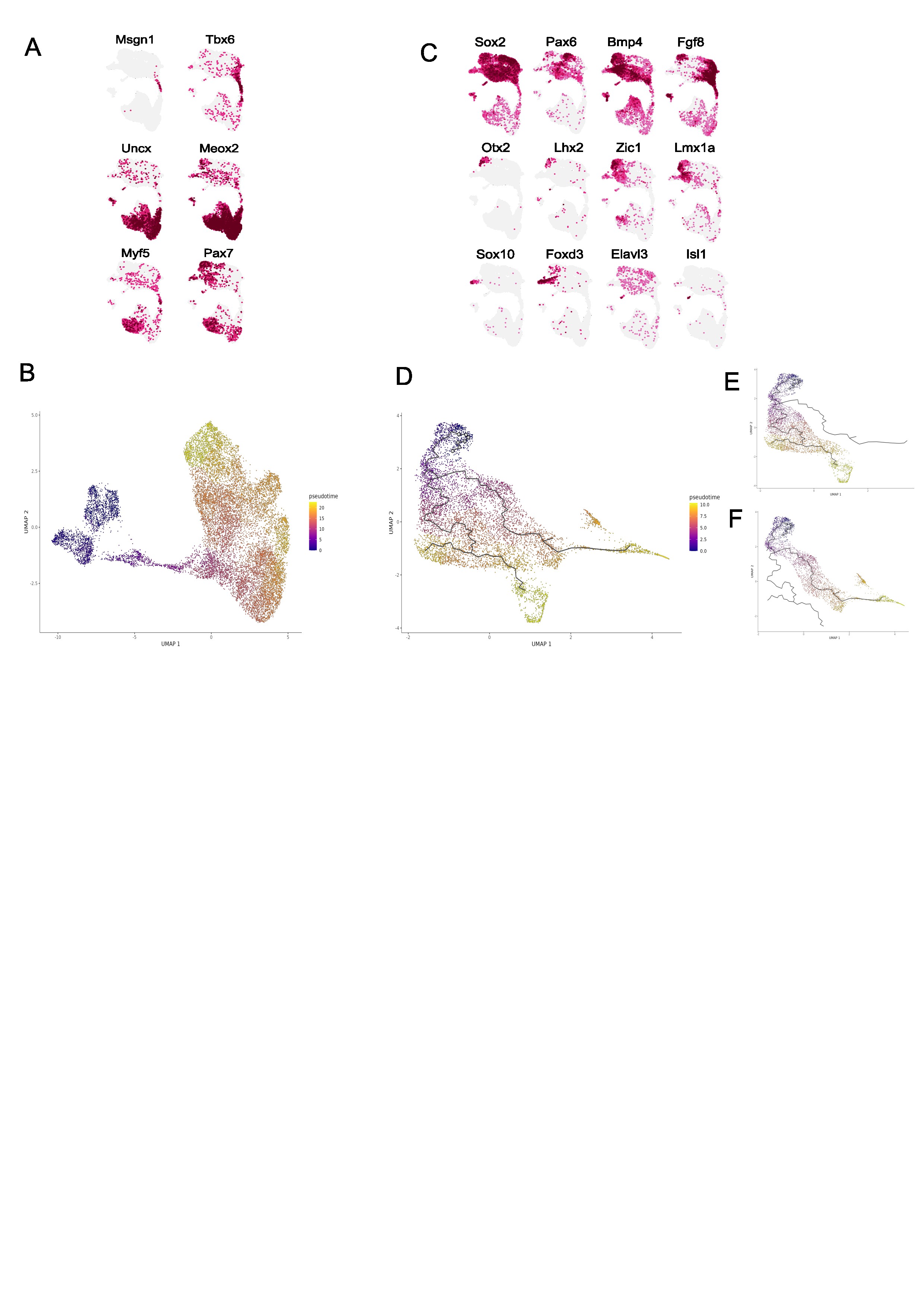
